# Targeted Deletion of Vitamin D receptor Gene in Mammalian Cells by CRISPR/Cas9 Systems

**DOI:** 10.1101/040147

**Authors:** Tao zhang, Ling Wang, Kun Xu, Chonghua Ren, Zhongtian Liu, Zhiying Zhang

## Abstract

CRISPR/Cas9 system has become a new versatile technology for genome engineering. It utilizes a single guide RNA (sgRNA) to recognize target sequences in genome function, and activates Cas9 endonucleases to cut the locus. In this study, we designed two target sites from conserved regions of vitamin D receptor (*VDR*) gene in mammalian cells, which cover more than 17 kb of chromosome region depending on the species. The efficacy of single sgRNA mediated gene specific modification was about 22% to 36%. Concurrently, targeted deletions of the intervening genomic segments were generated in chromosomes when the two sgRNAs worked simultaneously. The large genomic DNA segments ranging from 17.8Kb to 23.4 Kb could be precisely deleted in human and mouse chromosomes. Furthermore, the expression level of 24-hydroxylase (*CYP24A1*) regulated by VDR was significantly increased in cells treated with *VDR* CRISPR/Cas9 vectors. This study showed that CRISPR/Cas9 system can be employed to generate large genomic segment deletions in different species, providing sgRNAs are designed within conserved regions.

## Introduction

Vitamin D mediates a variety of biological functions such as calcium homeostasis, calcium reabsorption in the kidney, calcium mobilization in bone, cell differentiation and proliferation to many target tissues(DeLuca 2004). Most, if not all, the biological actions of vitamin D are believed to be exerted through the vitamin D receptor (VDR)-mediated control of target genes (Germain et al. 2006; Morrison et al. 1989). VDR is a member of the nuclear hormone receptor super-family of transcription factors that regulate gene expression in a ligand-dependent manner(Mangelsdorf et al. 1995). Mutations in the *VDR* cause the disease known as hereditary vitamin D resistant rickets (HVDRR) (Malloy et al. 1999). Through DNA microarray technology, 95 genes were identified that displayed different changes of expression level in *VDR* null mice, of which 28 genes were up-regulated and 67 were down-regulated (Li et al. 2003). Using whole body *VDR*^-/-^ mice, Claudin2 (*CLDN2*) gene had been demonstrated to be a direct target of the transcription factor VDR in cultured human intestinal epithelial cells(Zhang et al. 2015). *VDR* has been previously considered to regulate key steps in the hair cycle, and it has already been shown that mutations in *VDR* cause alopecia in humans and mice (Malloy et al. 2009). However, the complete profile of *VDR* action is still unknown, and precise targeted editing of *VDR* is critical to understanding the biological functions of *VDR*, which could be the key to development of novel therapeutic modalities for *VDR*-related diseases.

Targeted genomic editing is a powerful technology in revealing gene functions, gene therapy of human genetics for human diseases, generation of models and breeding animals with desired traits. Based on naturally occurring spontaneous homologous recombination, transgenic mice were generated via designed vectors with large homology arms (Norman 1995; Walters 1992). However, target efficiency was extremely low in the presence of targeted vector in other mammalian cells. Thus, this technology cannot be widely applied in other animals besides mice. It was illustrated that the introduction of DNA double-strand break (DSB) could trigger the efficiency of homologous recombination significantly in cells (Wyman and Kanaar 2006). Generally, endonucleases have the ability to generate DSB in specific DNA sequences. Nevertheless, the target site recognized by a natural endonuclease is not unique in genomic sequences. Thus, artificial nucleases were designed to cleave a specific DNA sequence, and generate a unique DSB in target cell genome. In recent years, several target genome-editing technologies have been developed and efficiently edited genomes in various types of cells and organisms. Zinc-finger nucleases (ZFNs) were the first generation artificial nuclease to be widely applied in insects, plants and animals (Remy et al. 2010). Another efficient genome targeting modification tool is transcription activator-like effector nucleases (TALENs), which offer far more attractive advantages in comparison to ZFNs, and have been rapidly and widely used to perform precise genome editing in a variety of organisms and cell types.

A novel genome editing platform based on clustered regularly interspaced short palindromic repeats(CRISPR)/CRISPR associated (Cas) protein system provides adaptive immunity against viruses and plasmids in bacteria and archae (Horvath and Barrangou 2010; Wiedenheft et al. 2012). The type II CRISPR/Cas9 of *Streptococcus pyogenes* is is a relatively simple CRISPR/Cas system, and only involves a single effector enzyme to cleave dsDNA. Given this advantage, it has rapidly been developed into a viable genome editing tool (Jinek et al. 2012). CRISPR/Cas9 nuclease is distinct from ZFNs and TALNEs, and it mediates genome editing following the rule of targeted DNA recognizing and cleavage by designed short guide RNAs (gRNA) recognizing target and endonuclease Cas9, respectively. Since the emergence of CRISPR/Cas9, scientists have devoted their efforts to promulgate the use of CRISPR/Cas9 system on the basis of facilitation of genome editing in mammalian cells. Zhang Feng achieved this goal, and developed a plasmid that contained both hspCas9 nuclease and a functional gRNA (Ran et al. 2013). Since then, the CRISPR/cas9 nuclease has become a dominant genome editing platform, and has been successfully used to generate target gene modified cells in plants and animals (Mali et al. 2013; Nemudryi et al. 2014; Bortesi and Fischer 2015; Tu et al. 2015).

In this study, we designed and constructed CRISPR/Cas9 nuclease to cut two target sites in the conserved sequences of *VDR*. The target sequences are exactly the same between humans and mice, and we aimed to use one target vector to achieve *VDR* targeted modification in 293T and C2C12 cell lines. Additionally, the target efficiency of CRISPR/Cas9 system on conserved sites was evaluated and compared in different cell types. This study displayed that CRISPR/ Cas9 system induced high rate mutations at two target sites of *VDR* in both mouse and human cell lines, and achieved large fragment deletion in respective chromosomes

## Materials and Methods

### Construction of CRISPR/Cas9 target vector and reporter vector

The sgRNA-Cas9 co-expression plasmid pX330-U6-chimeric-dBsaI-CBh-hspCas9 as parent vector was obtained from Addgene (http://www.addgene.org/), which harbors two different sticky ends by BsaI digestion (Cong et al. 2013). To construct VDR target vectors, the target dsDNA with sticky ends were generated via direct annealing of two oligonucleotides VDRT1F and VDRT1R (Table 1), in which sticky ends sequences exactly match BsaI ends in Cas9 parent vector. Thus, VDRT1 was cloned into the parent vector between two BsaI sites. Subsequently, sequencing was performed with U6F primer to confirm VDRT1 target vector, designated pX330-U6-VDRT1-CBh-hspCas9. Meanwhile, pX330-U6-VDRT2-CBh-hspCas9 as VDRT2 target vector was obtained with the same strategy.

**Table1.**
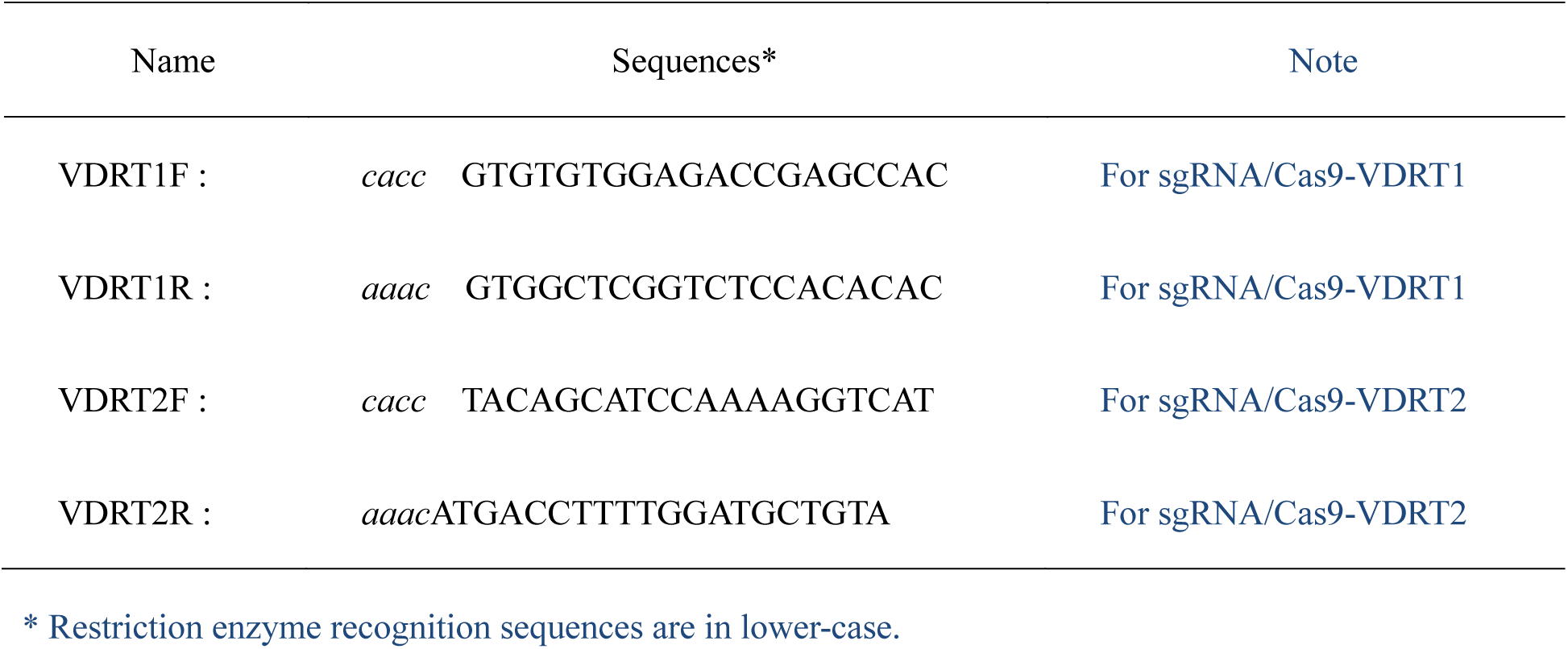
DNA oligos of sgRNA for CRISPR/Cas9 expression plasmids construction

pCAG-puro-NB-T2A-EGFP backbone plasmid was used to construct CRISPR/Cas9 reporter vector, in which puromycin resistant gene (Puro^R^) was separated by NotI and BamHI enzyme sites flanking with two 200bp direct repeats of *Puro*^*R*^. In order to insert *VDR* target sites into backbone plasmid, two oligonucleotides for each target were designed and synthesized **(Table 2)**, and target DNA fragments harboring PAM sequence and NotI and BamHI sticky ends were generated by direct annealing. Then *VDR* target fragments were cloned into pCAG-puro-NB-T2A-EGFP between NotI and BamHI sites to achieve two reporter plasmids, designated pCAG-puro-VDRT1-T2A-EGFP and pCAG-puro-VDRT2-T2A-EGFP, respectively.

**Table2.**
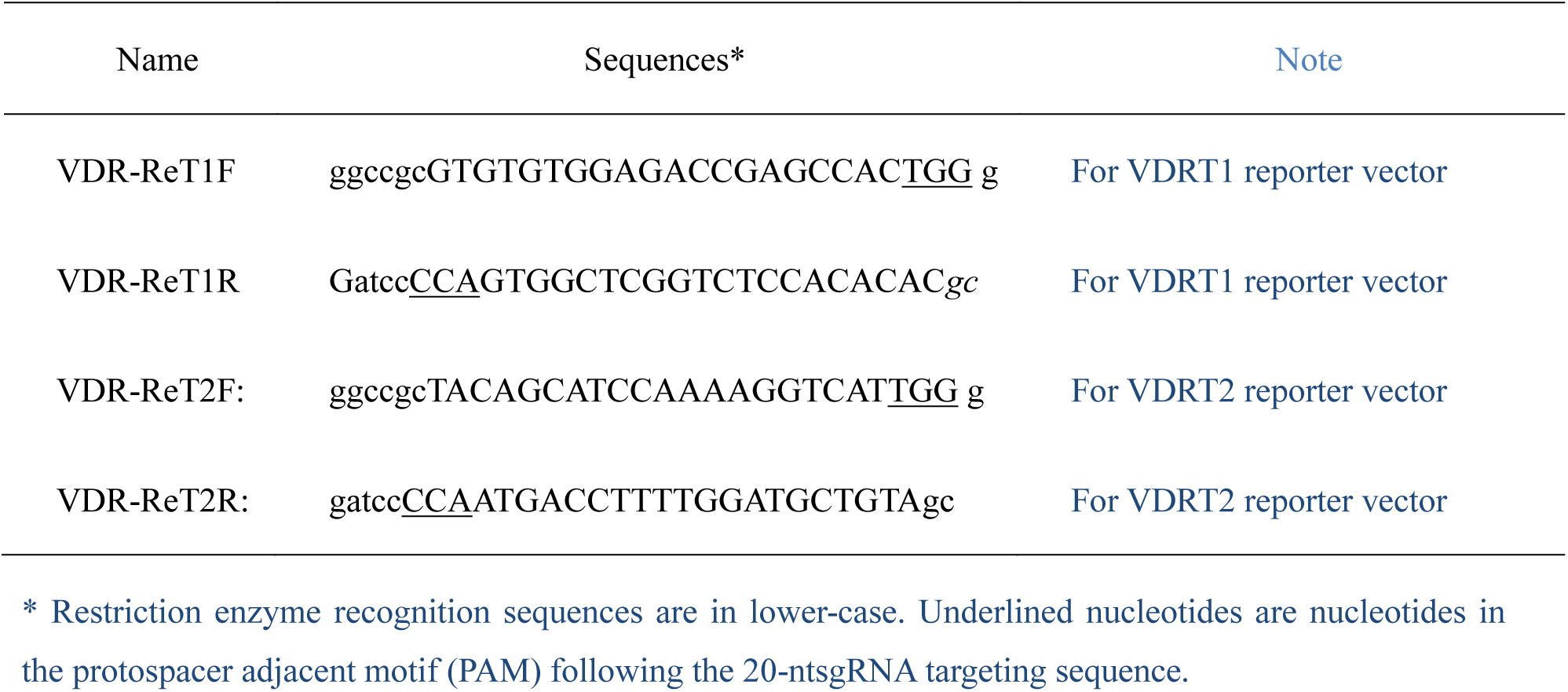
DNA oligos of target site for report vector construction

### Cell culture and transfection

HEK293T (human embryonic kidney cell line) and C2C12 (mouse myoblast cell line) cells were obtained from the American Tissue Collection Center (ATCC). These two cell lines were cultured in DMEM (Dulbecco’s modified Eagle’s medium; Gibco) supplemented with 10% fetal bovine serum and 1% penicillin/streptomycin, and were maintained at 37°C and 5%CO_2_. The 293T and C2C12 cells were transfected using NeoFect™ DNA transfection reagent (Neofect biotech, Beijing). According to manufacturer’s instructions, 2μg Cas9 expression vector and 1μg report vector were added into each cell culture of 6-well plates. At 48 hour post-transfection, puromycin enrichment was launched to enrich cells containing restored *puro*^*R*^ in the reporter vector. After 48 hours for puromycin treatment, cells were maintained in a fresh medium without puromycin for 24 hours, and then the genomic DNA was extracted for PCR.

### Genomic DNA isolation and PCR detection

Total genomic DNA was extracted from human 293T cell and mouse C2C12 cell according to the phenol-chloroform procedure (Malumbres et al. 1997). The target and off target regions were amplified by PCR with Taq DNA polymerase (Fermentas Inc) according to the manufacturer’s instructions. Using 50-100ng of genomic DNA as template, the cycling program was 95 **°C** for 5 minutes followed by 35 cycles of 94 **°C** for 30 seconds, 55 **°C** for 30 seconds, and finally 72**°C** for 30 seconds. Primers used for PCR were listed in table 3.

**Table3.**
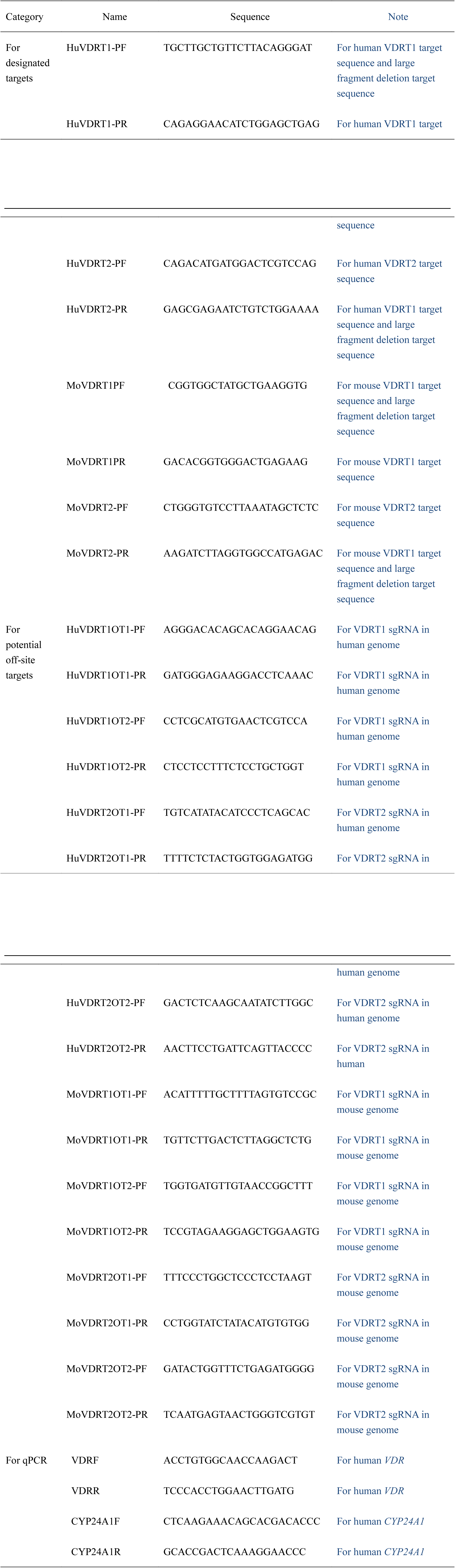
PCR primers for target and off target sequences amplification

### T7 Endonuclease I (T7E1) assay

T7E1 assay was performed as previously described (Kim et al. 2009; Ran et al. 2013). PCR products were briefly purified using a DNA purification kit (Hangzhou Bioer Technology Co.,Ltd.). Subsequently, purified products were denatured by heating (for 2 min at 94°C) and annealing (94°C to 85°C at 2°C per second,85°C to 25°C at 0.1°C per second), followed by digestion with the mismatch-sensitive T7 endonuclease I **(NEB)** and then analyzed using 2% agarose gel electrophoresis.

### Deletion frequencies of large fragment by digital PCR

Genomic DNA of target cells were serially gradient diluted in double distilled water to the gradient concentrations of 100ng, 33ng, 10ng, 3.3ng, 1ng, 330pg, 100pg, 33pg, 10pg, 3.3pg and 1pg. Theoretically, 3.3pg ([3.0 × 10^9^ bp × 650 g/mol/bp]/6.0 × 10^23^) of genomic DNA per reaction was considered to be equivalent to “a haploid genome”.(He et al. 2015; Flores et al. 2007) Control PCR was carried out by using the primers of wild-type genes to confirm this gave target PCR products at a dilution containing 3.3pg DNA reaction. In fact, the target PCR product rose at a dilution containing 10 pg of DNA per reaction, and no PCR product was observed at the concentration of 3.3pg of DNA. To calculate the deletion frequencies, 11 wild-type gene reactions and 11 target fragment reactions were performed by related primers in parallel at each dilution point, respectively. The lowest concentration of genomic DNA that gave rise to PCR products was determined and used to estimate deletion frequencies.

### RNA isolation and Real-time RT-PCR

Total RNA was extracted from target cells via Trizol reagent (TaKaRa, Dalian China). The first-strand cDNA was generated using a reverse transcription kit (TaKaRa) with random primers. Real-time quantitative PCR was performed in triplicate samples using a SYBR green kit (Invitrogen, Thermo Fisher) on the AB Step one plus system. Human *GAPDH was* taken as the reference gene. The 2-ΔΔCT algorithm was employed to estimate the relative expression level of each gene. The sequences of primers were listed in Table 3.

### Target sequencing and sequence alignment

The purified PCR products of target DNA were cloned into the T-vector using the pGEM-T Kit (Promega). For each target site, 10 independent colonies were chosen for sequencing detection using the T7F primer by ABI 3130 automated sequencer (**BeijingAuGCTCo.,Ltd**). DNA sequence alignment was performed to compare *VDR* target locus with wild-type sequence.

## Results

### Design and construction of CRISPR/Cas9 system for *VDR* editing

*VDR* is located on human chromosome 12 and mouse chromosome 15, respectively. To achieve target knockout *VDR* in human and mouse cells, *VDR* sequences of humans and mice were analyzed to choose two target sites in the conserved region in both human and mouse genomes. According to the design principle and program of CRISPR/cas9 (http://crispr.mit.edu/, Zhang Feng Lab), we designed two target sgRNAs (VDRT1 and VDRT2), which respectively target exon 4 and exon 7 of *VDR* in human genome and exon 3 and exon 7 of the mouse orthogonal gene. The two target sites are separated by 23.4kb DNA fragment in human genome and 17.8 kb in mouse genome (Fig.1A). Therefore, we constructed sgRNA and Cas9 protein co-expression vectors pX330-U6-VDRT1-CBh-hspCas9 and pX330-U6-VDRT2-CBh-hspCas9 for targeting VDR gene in both human and mouse culture cells.

**Fig.1.**
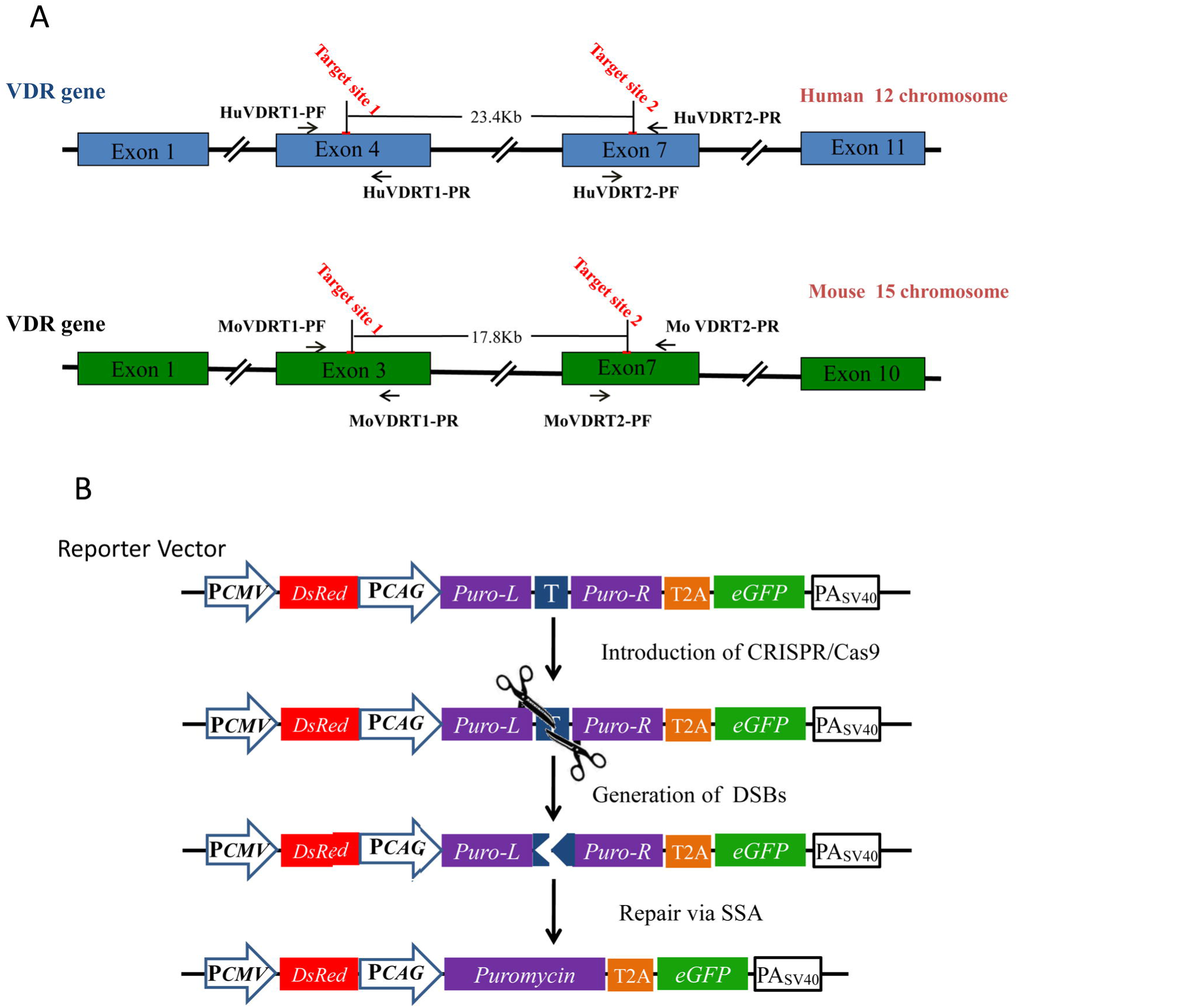
Strategy for target editing in human and mouse chromosome and target cells enrich systems. (A) Schematic diagram of location of two *VDR* targeting sites in human and mouse chromosomes. The red lines stand for the locations of two target sites in *VDR*, and the distances between the two target sites and each sequence of target sites are shown. The primers for the PCR amplified target sites sequence are indicated by a black arrow. (B) Schematic diagram of reporter vector characteristics. DsRed gene was a reporter gene to test the transfection efficiency of CRISPR/Cas9 system. During the introduction of CRISPR/Cas9, the disrupted puromycin resistance gene (Puro^R^) was repaired by single strand annealing (SSA), resulting in restored PuroR and eGFP.

In order to enrich genetically modified cells, we designed and established a screening system (Fig.1B) based on reporter plasmid that could be used to test the transfection efficiency of CRISPR/Cas9 target vectors and enrich target cells simultaneously. In this reporter vector, DsRed gene was used to detect the transfection efficiency, and puromycin resistant gene (Puro^R^) and eGFP were used as reporter genes to validate cleavage activity and enrich positive cells. To simulate target sites in genomes, Puro^R^ was separated by the target DNA fragment with PAM sequences flanking two 200bp direct repeats, and therefore, this insertion disrupts its open reading frame. Once designed CRISPR/Cas9 cut the target sequence in the reporter plasmid, the Puro^R^ gene was repaired by SSA-mediated DNA repair mechanism to restore wild-type Puro^R^, and provided the target cells the ability to survive under puromycin selection pressure in medium. Conversely, cells without restored Puro^R^ gene were unable to survive in puromycin medium. Therefore, cells with *VDR* targeted-modification were enriched.

### Cleavage efficiency of each VDRT sgRNA in HEK293T and C2C12cells

In order to test the cleavage efficiency of CRISPR/Cas9 on target sites in conserved sequence of *VDR*, CRISPR/Cas9 expression vectors and their corresponding report plasmids were co-transfected into human 293T cells. After transfection 24h, Red fluorescence and green fluorescence positive cells were observed (Fig. 2A). Meanwhile, puromycin was added into the medium for screening and enrichment of positive clones for 48h, and cells were harvested to extract total genome DNA for further analysis. CRISPR/Cas9-induced *VDR* indels were measured using T7 endonuclease I (T7E1), which cleaves heteroduplexes formed by the hybridization of mutant and wild-type *VDR* target sequences or two different mutant sequences. The mutation frequencies of the two *VDR* sites in 293T cells were 36% and 31%, respectively (Fig. 2B). In addition, the mutation frequencies of the VT1 and VT2 sites in enriched C2C12 cells were 26% and 22% by T7E1 assay.

**Fig.2.**
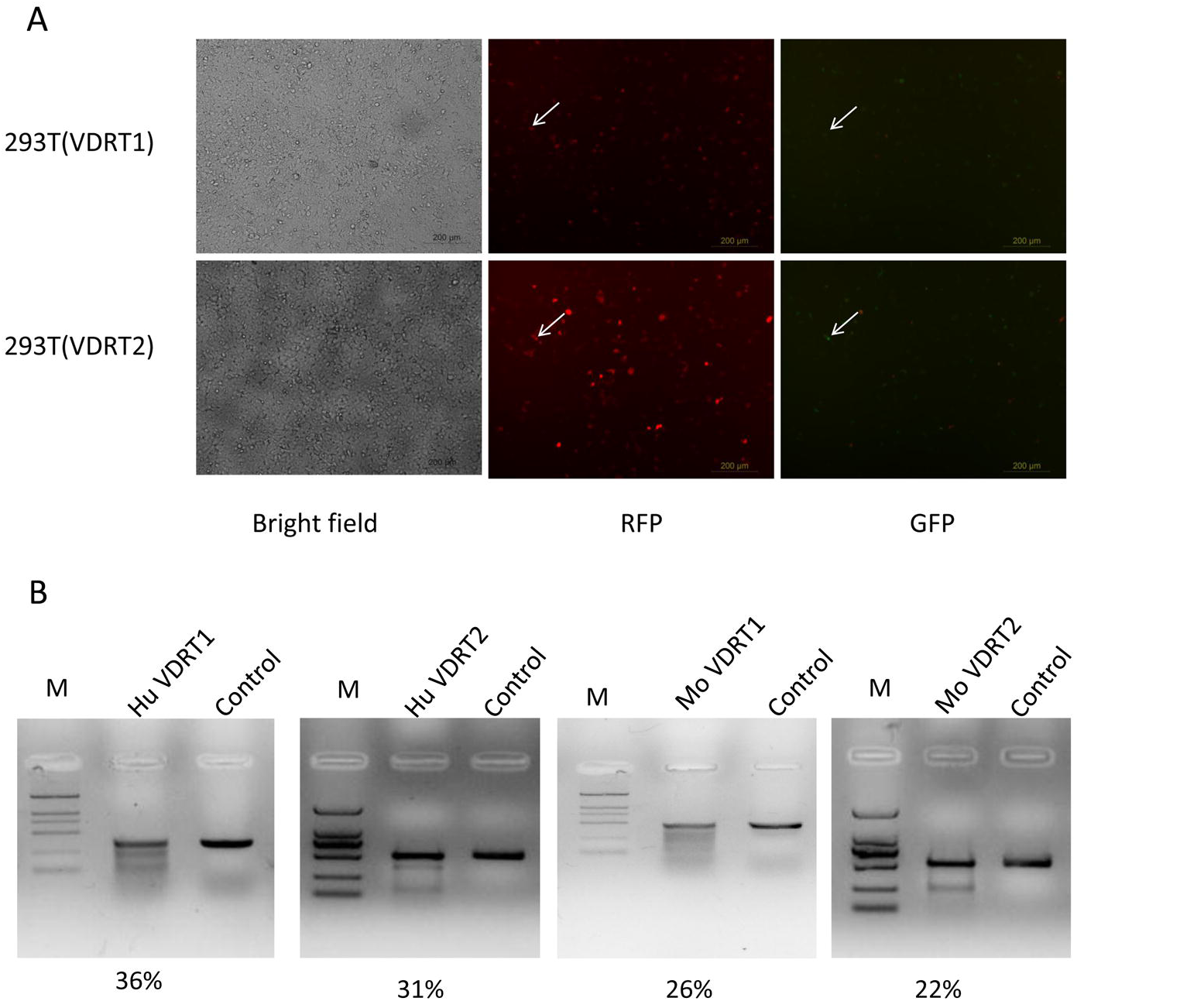
Knockout of *VDR* by CRISPR/Cas9 in human 293T cells and mouse C2C12cells. (A) The transfection efficiency and activity of CRISPR/Cas9 in HEK293T cells. (B) The efficiency of CRISPR/Cas9 mediated cleavage at two target sites in HEK293T cells and C2C12 cells.

To further confirm the mutation frequency induced via CRISPR/Cas9, we cloned the PCR product surrounding the target sites amplified from these enriched cells. Ten clones from each human VDRT1 and VDRT2 sites were randomly picked for direct DNA sequencing. The sequencing results demonstrated that random indels were detected in 4 colonies of VDRT1 and 2 colonies of VDRT2, respectively (Fig.3A and B). Deletion of 9nt was observed in two independent colonies of VDRT1, though 7nt deletion was verified in only 1 colony. Additionally, 44nt insertion was also found in VDRT1. By comparison, only deletions of 1 nt and 4nt were detected in VDRT2 site. However, mutations from twenty clones were not observed at VDRT1 and VDRT2 in mouse C2C12 cells.

**Fig.3.**
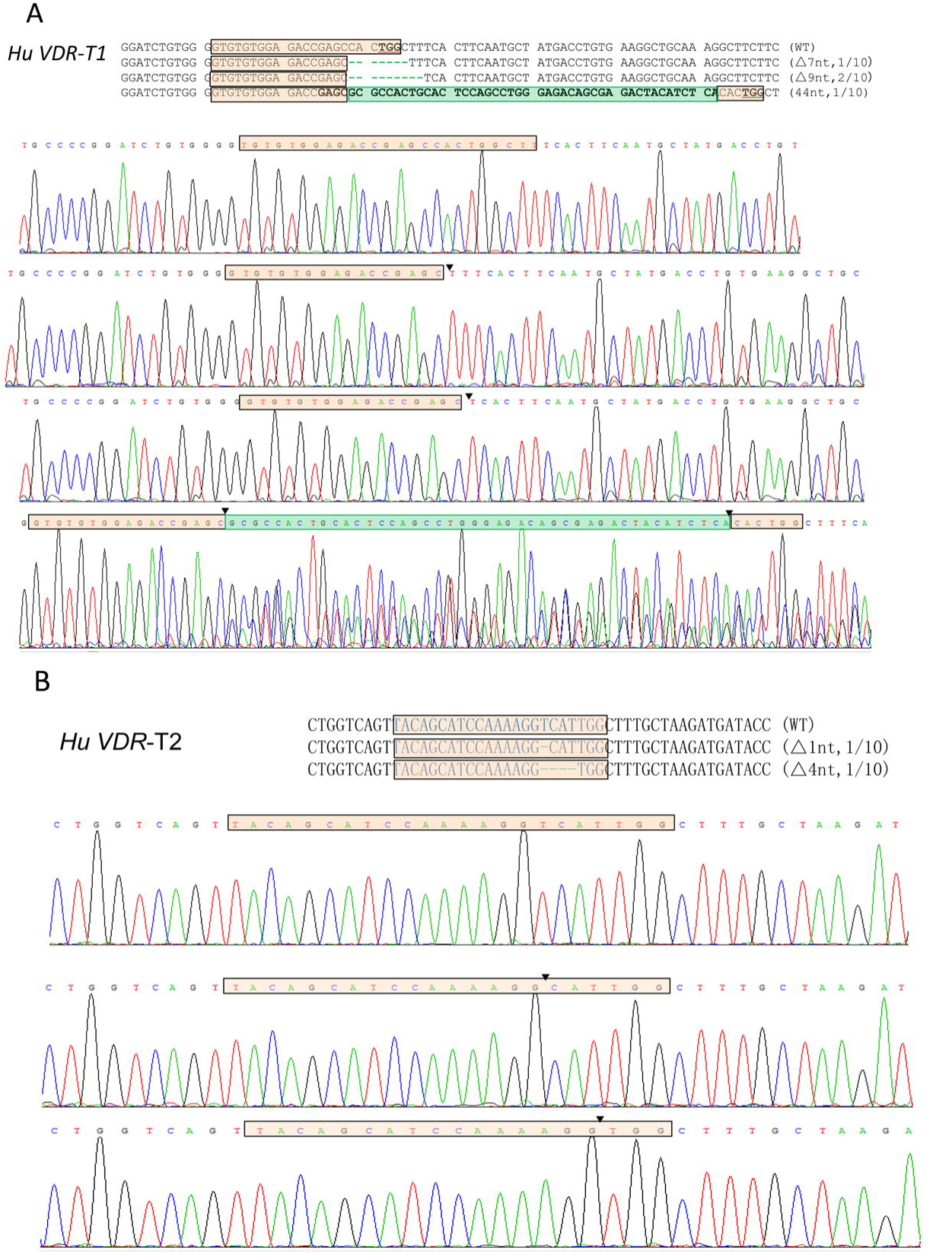
CRISPR/Cas9 mediated *VDR* targeted editing in human 293T cells. (A) Targeted indel mutations via VDRT1 CRISPR/Cas9. (B) Targeted indel mutations by VDRT2 CRISPR/Cas9. Target sequences were highlighted with pink boxes, and PAM sequences were highlighted as underlined. Green boxes stand for insertion sequences. (▼, deletion junction; Δ,deletion; + insertion)

### Highly efficient large fragment deletion in *VDR* using CRISPR/Cas9

After verifying each single sgRNA activity on VDRT1 and VDRT2 target sites, we contemplated whether we could delete the large chromosome segment between VDRT1 and VDRT2 sites by co-transfecting plasmids expressing two sgRNAs in addition to Cas9. We examined the efficiency of deleting the large chromosome segment in both human and mouse cells. One pair of primers, VDRT1PF and VDRT2PF, which are located upstream of the VDRT1 site and downstream of the VDRT2 site respectively, was used to amplify *VDR* chromosome DNA. If a large DNA fragment had been deleted from the chromosome, a 500bp DNA fragment was amplified. Otherwise, no product could be amplified when wild-type chromosomes were used as templates. After being enriched with puromycin for 72h, the transfected cells were harvested and the genomic DNA was prepared for PCR amplification. As shown in Fig 4A, we detected roughly 500bp PCR products using primers VDRT1PF and VDRT2PF from Cas9 treated cell genome DNA, and no products were obtained using wild-type cell genomes as templates (Fig.4A).

**Fig.4.**
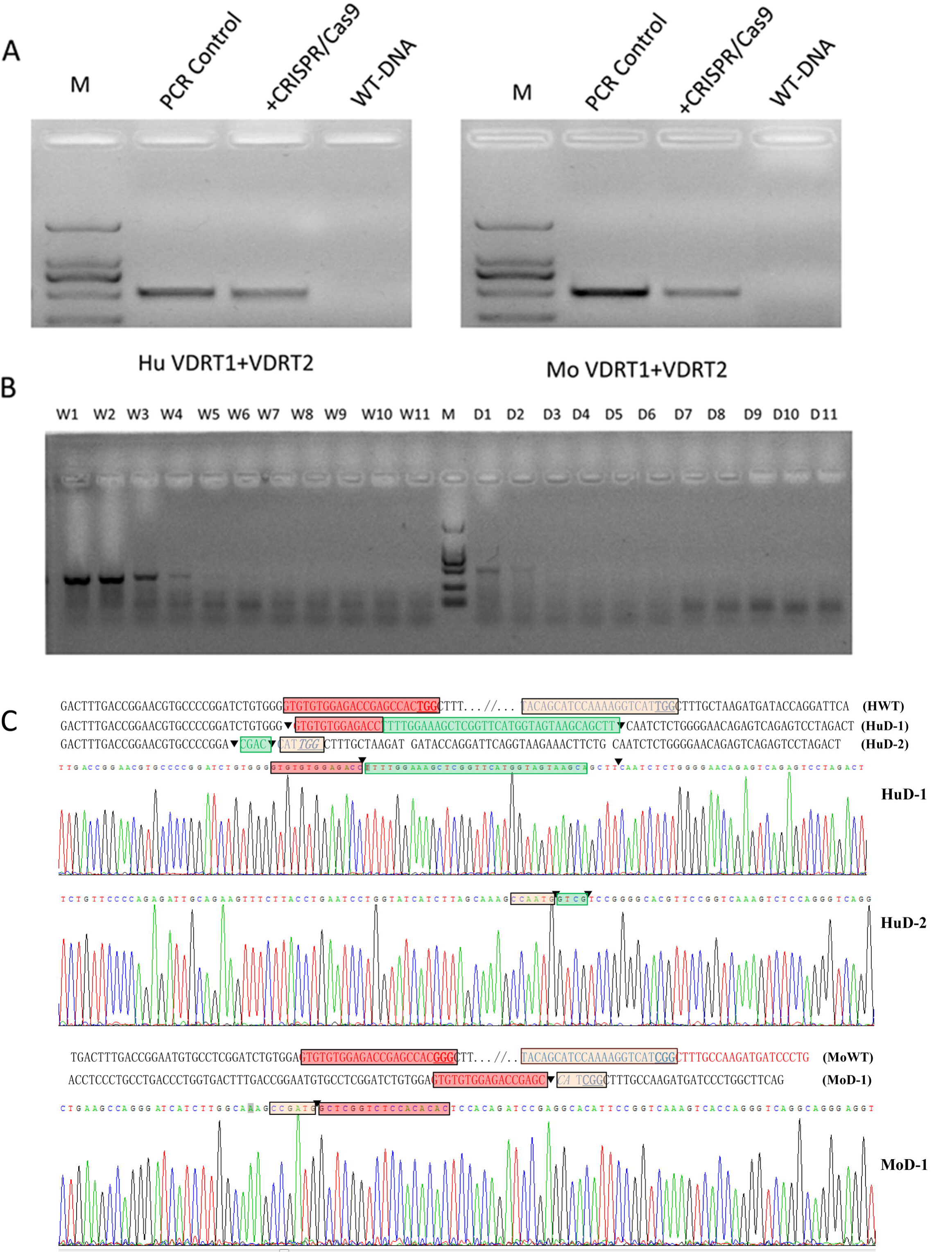
CRISPR/Cas9 mediated large fragment deletion within *VDR* in HEK293T cells and mouse C2C12 cells. (A) The large fragment deletion was identified by PCR. The PCR products of lane 3 (+CRISPR/Cas9) and lane 4 (WT-DNA) were amplified from treated cells and wild type genomic DNA, respectively. The non-relative PCR was performed to confirm PCR system normal in lane 2. (B) “Digital” PCR products identifying the larger genomic deletions within the *VDR*. The strong 400bp band in all lanes W1-W11 (on Mark right) is a non-specific amplification product. The 500bp bands from D1-D11 (on Mark right) correspond to the expected size of the PCR product in the event of large fragment deletion of the intervening sequence. (C) The indel mutations were identified by Sanger sequence. Target sequences were highlighted with red boxes and pink boxes, and PAM sequences were highlighted as underlined. Green boxes stand for insertion sequences. (▼, deletion junction)

Furthermore, deletion efficiency was measured via gradient dilution assay. In this assay, gradient dilution genomic DNA samples were used as a template for PCR to detect the target fragment and a wild-type gene. PCR products from each dilution point would be revealed by gel electrophoresis. In this study, the positive PCR products of wild-type gene were detected up to the concentration of 3.3ng genome DNA reaction, and the target fragments of large fragment deletion *VDR* gave positive PCRs up to the dilution containing 33ng of DNA (Fig.4B). Therefore, the frequencies of large fragment deletions reached 10% at *VDR* locus in human cells. Subsequently, PCR products sequencing results confirmed a 23.4Kb deletion in human and 17.8Kb deletions in mouse chromosomes, respectively. Compared with wild-type sequences, truncated *VDR* target sites and random indels were observed between VDR1 and VDR2 target sites in both human and mouse chromosomes (Fig.4 C).

### Expression levels of *VDR* and *Cyp24A1* in the target cells

We analyzed the expression level of *VDR* and 24-hydroxylase (*Cyp24A1*) to detect the effect of *VDR* target deletion in 293T cells. The *CYP24A1* enzyme catalyzes the first step in the catabolic pathway, converting 1α,25 (OH)_2_D3 into the less active intermediate 1,24,25(OH)_3_D3(Haussler et al. 1998). The *CYP24A1* gene is significantly up-regulated by 1α, 25(OH)_2_D3 through two VDRE in the proximal promoter region (Armbrecht et al. 1998; Chen and DeLuca 1995). The results show that the mRNA level of VDR was significantly increased in cells treated with T1 and T2 sgRNA compared with the control, and the expression level of *VDR* decreased in cells transfected with both T1 and T2 sgRNAs. However, the mRNA levels of *CYP24A1* significantly decreased in all groups treated with VDR sgRNA compared with the control group (Fig.5). The results demonstrated that CRISPR/Cas9 modified *VDR* could impact its relative gene expression in mammalian cells.

**Fig.5.**
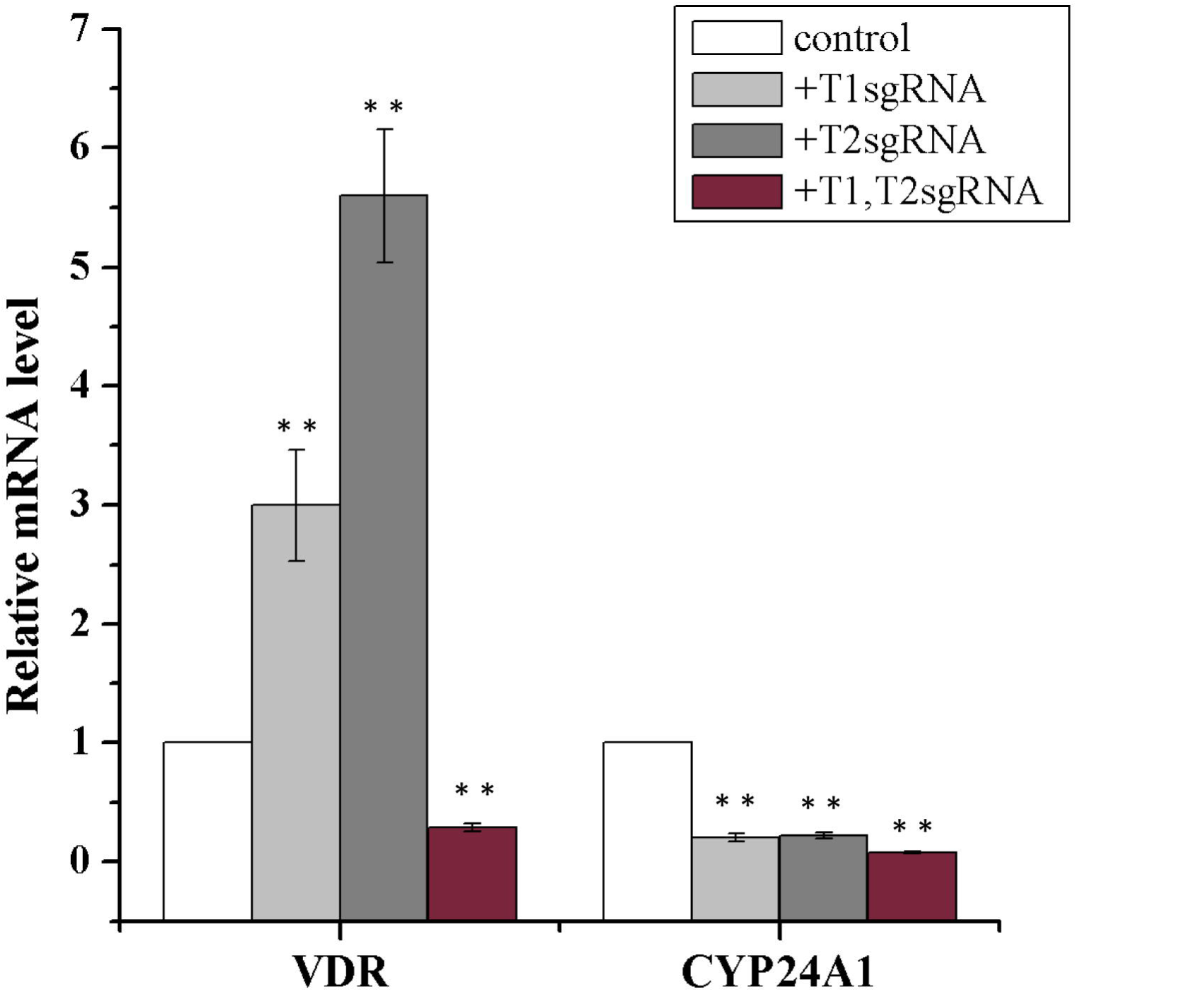
Expression levels of *VDR* and *Cyp24A1* different treated cells. The control group indicated that the cells were treated with no related sgRNA. T1 sgRNA, T2 sgRNA show that the cells were treated with CRISPR/Cass9 vectors of VDRT1 and VDRT2, respectively. And T1, T2 sgRNA stands for cells co-transfected with VDRT1 and VDRT2.

### Off-targeting detection of CRISPR/cas9 in human 293T cell and mouse C2C12

CRISPR/Cas9 system has become a newly developed, powerful tool for targeted genome editing. However, high target efficiency was accompanied by high off-targeting effects, a finding consistent with reports from other studies. Compared with ZFNs and TALENs, the rate of off-target cleavage via CRISPR/Cas9 could be higher, because only 20bp recognition sequence of each CRISPR/Cas9 is shorter than target sequences of a pair of ZFNs and TALENs. Many researches revealed that CRISPR/Cas9 had unexpected off-target effects in culture cells and several organisms (Fu et al. 2013; Pattanayak et al. 2013; Cradick et al. 2013). In addition, the off-target sites were much more similar with target sites, when target DNA sequences contained insertions (‘DNA bulge’) or deletions (‘RNA bulge’) compared to the RNA guide strand, and these genomic sites could be cleaved by CRISPR/Cas9 systems(Lin et al. 2014). In order to evaluate off-targeting effects of these CRISPR/Cas9 nucleases, we chose two candidate loci for each target site with high potential cleavage in human and mouse genomes via the program of CRISPR/cas9 (http://crisp potential off-targetr.mit.edu/, Zhang Feng Lab) [Table 4]. Through T7E1 assay, the results showed the pX330-U6-VDRT1-CBh-hspCas9 plasmid induced off-target mutations at two potential off-target sites HVOT1 and HVOT2 with frequencies 8% and 6% in HEK293T cells. No mutations were detected in mouse C2C12 cells however. In addition, no obvious off-target effect was detected from pX330-U6-VDRT2-CBh-hspCas9 at the other four off-target sites in human or mouse genomes (Fig. 6). These results suggest that CRISPR/Cas9 could induce mutations at some sites in chromosomes, but the specific sgRNA could avoid or reduce the off-target effect in genome editing research.

**Table4.**
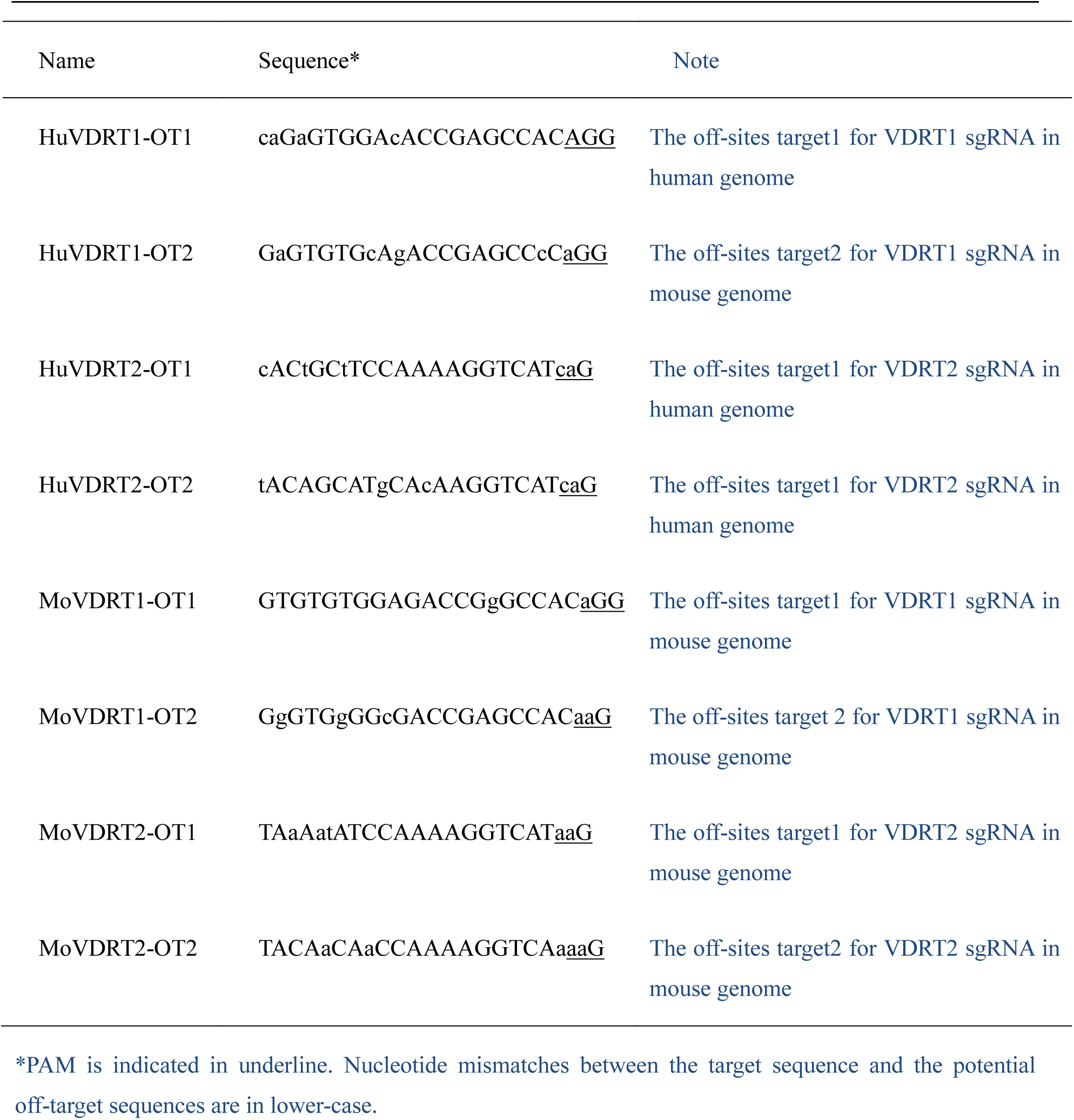
The sequences of potential 1 off-target sites in human and mouse

**Fig.6.**
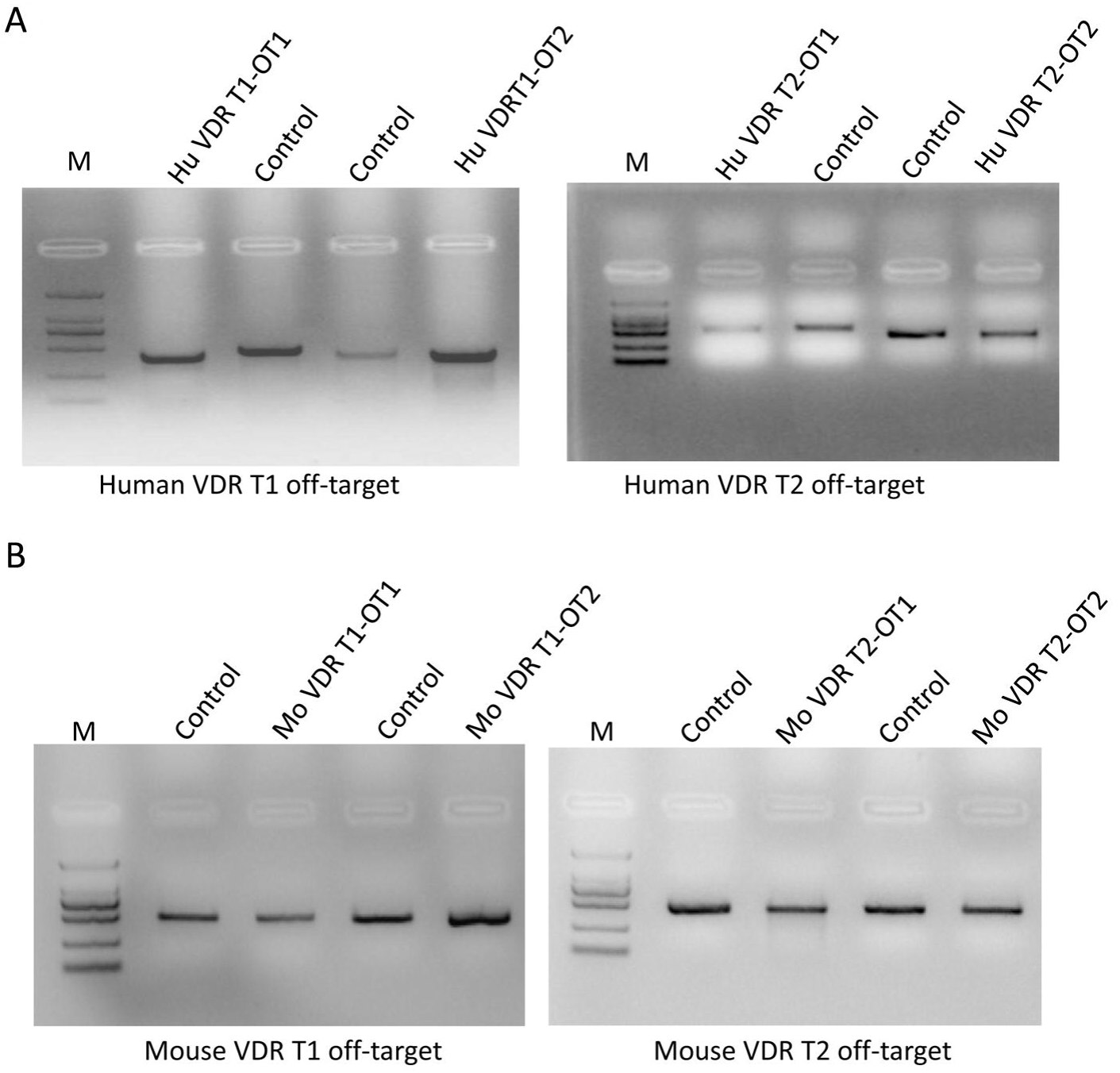
Detection of CRISPR/Cas9 mediated off-targeting. (A) The off target effect of VDR sgRNA1 and sgRNA2 in HEK293T cells. (B) The off target effect of VDR sgRNA1 and sgRNA2 in C2C12 cells.

## Discussion

CRISPR/Cas9 nucleases are powerful tools for precise genomic editing in various species, which have greatly simplified the process of constructing target vectors and have simultaneously enabled low-cost access of this system to the entire field of biomedical research. In our study, we presented the CRISPR/Cas9 system effectively modified conserved sequences in human and mouse *VDR* with relatively high efficiency. *VDR* specific sites editing cell lines or model animals could be widely used in VDR biologic function exploring, *VDR* relative diseases research and novel medication development. Hereby, we engineered two CRISPR/Cas9 nucleases, which targeted VDRT1 and VDRT2 sites and successfully modified VDR in both human and mouse cells. To achieve much more nuclease-targeted cells, a dual-gene report system was employed to screen and enrich genetically modified cells.

Except PAM sequences, target sites in human and mouse genomes are identical. By comparison, it is easy to understand that the efficiency of CRISPR/Cas9 varied greatly between VDRT1 and VDRT2 sites in HEK293T cells as well as C2C12 cells. CRISPR/Cas9 induced VDRT1 mutation efficiency via NHEJ in 293T cell was 36%, which was much higher than the mutation rate in C2C12 cells (26%). Meanwhile, CRISPR/Cas9 mediated VDRT2 modified efficiency was 30% and 21% in human and mouse cells, respectively. Obviously, these nucleases worked far more efficiently in human 293T cells than mouse C2C12 cells. We speculated that different positions of *VDR* target sites in human and mouse chromosome might affect CRISPR/Cas9 cleavage activities. Thus, these results suggest that chromosomal structure might play an important role in regulation of CRISPR/Cas9 activity.

Another reasonable factor for this phenomenon is the difference of PAM sequences for identical sites in HEK293T and C2C12 cell lines, in which TGG as PAM sequence for VDRT1 and VDRT2 in human genome, and GGG and CGG PAM sequences for VDRT1 and VDRT2 in mouse genome. Though we cannot confidently conclude that the cleavage effect mediated by TGG PAM sequence was higher than GGG or CGG, our results indicate that PAM sequences may play a critical role in the activity of CRISPR/Cas9 system.

In this study, there were any mutant alleles in 20 clones of mouse VDR target, but the results of the T7E1 assay indicate that VDR alleles were mutated in cultured mouse cells. We thought that three reasons should been consider: firstly, the plasmid transfection efficiency was very low in mouse cells, and the positive mouse cells were very limited by enrichment. Secondly, T7E1 assay is a direct and coarse approach in mutation detection, and the detecting results may be inconsistent with the sequencing. Another, the “flanking” sequences of target sites may impact the result from T7E1 assay.

Taken the advantage of multiple sites target editing, CRISPR/Cas9 systems greatly facilitate large DNA fragment deletion in chromosomes (Zhou et al. 2014b). Hereby, CRISRP/Cas9 nucleases targeting VDRT1 and VDRT2 were introduced into HEK293 and C2C12 cells together to achieve large fragments deletion in 293T cells and C2C12 cells. And fragments of 23.4Kb and 17.8Kb between VDRT1 and VDRT2 were deleted in human and mouse chromosomes, respectively. The efficiency of 23.4 Kb DNA segment deletion in 293T cells was up high to 10%, which was much higher than deletion of chromosomes in *Caenorhabditis elegans* (Chen et al. 2014) and rice (Zhou et al. 2014a). By comparison, another report demonstrated that the efficiency of 65 Kb large DNA fragment deletion was as high as 11.8% in mouse embryonic stem cells via microinjection of CRISPR/Cas9 expression plasmids. According to sequencing results, the breakpoint junctions indicated that CRISPR/Cas9 mediated DSBs were repaired by NHEJ mechanism, a noticeable disparity compared with the NHEJ repairing DSBs induced by ZFN and TALENs (Kim et al. 2013). It is likely that Cas9 has the characteristic of cleavage DNA between the third and fourth base pairs in the upstream of the PAM and generates blunt ends (Jinek et al. 2012), and the ligation of two blunt DNA ends may not require a micro-homology alignment process.

Off-target mutations and chromosomal translocations could lead to adverse biological effects, such as gene mutation, inactivation of tumor suppressors, and activation of oncogenes. Previous studies reported that high off-target effects were detected in the whole genome, especially in cultured cell lines. Recently, several groups revealed that off-target cleavage of CRISPR/Cas9 occurred at some genomic sites that differ with up to five nucleotides from the target sites (Pattanayak et al. 2013). In this study, eight potential off-target loci were selected to detect off-target efficiency. At two potential off-target sites HVOT1 and HVOT2, we detected off-target mutations at frequencies 8% and 6% via T7E1 assay in HEK293T cells, respectively. At the other six potential off-target sites, no mutations were detected in human or mouse genomes (Fig.6). Due to the poor sensitivity of T7E1 assay, it cannot detect off-target mutations that occur at frequencies <1%(Kim et al. 2009). By contrast, deep sequencing can measure off-target mutations that occur at frequencies ranging from 0.01% to 0.1%. Thus, the six potential off-target sites should be subject to further evaluation via more suitable methods.

To further analyze the off-target sequences, the two sites with obvious off target effect all possess AGG PAM motif, and these mismatches were not in the ‘seed sequence’. For the other six potential off-target sites, some sites have no NGG PAM while some mismatches were located in the ‘seed sequence’. Previous studies reported that NGG (or CCN on the complementary strand) sequences were sufficient for Cas9 targeting and that NGG to NAG or NNGGN mutations in the editing template should be avoided (Jiang et al. 2013). For protospacer, the point mutations within the ‘seed sequence’ (the 8 to 10 protospacer nucleotides immediately adjacent to the PAM) could abolish CRISPR targeting activity. The distal (from the PAM) positions of the protospacer (12 to 20) could tolerate most mutations (Semenova et al. 2011; Jinek et al. 2012; Wiedenheft et al. 2011). Thus, our results are consistent with previous reports, and confirmed this conclusion.

For researchers, the appropriate method to reduce off-target effect is the key issue and top concern in the application of CRISPR/Cas9 system. Scientists have developed various strategies to construct CRISPR/Cas9 system with low off-target effect. One is to optimize the CRISPR/Cas9, while the other is to choose specific target sites. In addition, Cas9 was modified to generate nickases, which introduce single-strand breaks in target sites as well as off-target loci. Subsequently, single-strand breaks in off-target loci are repaired without any mutations (Cho et al. 2014). A report indicated that RGEN ribonucleoproteins (RNPs) could greatly reduce off-target mutations by delivery of purified recombinant Cas9 protein and guide RNA into cultured human cells (Kim et al. 2014). Furthermore, software has been developed for assessing the off-target effect in genome editing by CRISPR/Cas9, which is a critical parameter for screening target sites (Cradick et al. 2014; Xie et al. 2014; Naito et al. 2015; Xiao et al. 2014).

One of the main goals of targeted gene modification is to achieve an ideal model for revealing gene functions. In this study, *VDR* expression levels were significantly different among these groups, but the expression of *Cyp24A1* decreased efficiently in cells treated with either a single or double sgRNAs. This phenomenon indicated that CRISPR/Cas9 mediated mutations of *VDR* diminished its function *in vivo*, which could provide an alternative model for VDR function research. We confused that the levels of *VDR* transcript were increased in cells transfected with T1 or T2 sgRNA constructs, but decreased in cells co-transfected with both T1 and T2 sgRNA constructs. We speculate that single site mutation in *VDR* disrupted the bio-active functions, but an unknown feedback system drove the cells to express more *VDR* mRNA. Therefore, levels of *VDR* transcript were increased in cells transfected with T1 or T2 sgRNA constructs. For large fragment deletion cells, the normal expression of *VDR* was disrupted seriously, thus levels of *VDR* transcript were decreased in cells co-transfected with both T1 and T2 sgRNA constructs.

In summary, CRISPR/Cas9 is an effective genome-editing tool for precise genomic modification. We constructed CRISPR/Cas9 systems and reporter vectors targeting conserved sequences of *VDR* in HEK293T cell and C2C12 cell lines. Specific modifications and large DNA fragment deletion were obtained after the introduction of CRISPR/Cas9 systems. This study provides a new technology platform for precise gene modification in conserved regions in different species.

## Acknowledgments

We wish to thank all of the colleagues in Professor Zhang’s lab for their excellent technical assistance and careful reading of this manuscript. This research was supported by the National Nature Science Foundation of China. This research was supported by grants from National Natural Science Foundation of China (NSFC)[31171186, 31402071], National Science and Technology Major Project of China [2014ZX0801009B] and Natural Science Basic Research Plan in Shaanxi Province of China [2013JQ3009].

